# Fire limits soil microbial dispersal and differentially impacts bacterial and fungal communities

**DOI:** 10.1101/2025.05.02.651892

**Authors:** Jacob R. Hopkins, Giuseppina Vizzari, Alison E. Bennett, Antonino Malacrinò

## Abstract

Fire is a globally pervasive force reshaping ecosystems, yet its influence on the ecological processes structuring soil microbiomes remains poorly understood. Using a meta-analysis of >2,600 amplicon sequencing samples across 19 global studies, we tested whether fire alters soil microbiome assembly processes, diversity, and ecological selection for pyrophilic specialists. Contrary to prevailing assumptions, we found that fire did not significantly shift ecological selection processes in bacteria or fungi but instead constrained dispersal, particularly reducing dispersal in bacterial and fungal communities, and increasing ecological drift in fungi. Despite limited evidence for ecological selection, fire consistently filtered for specialist taxa, increasing their relative abundance across microbial communities. Fire also reduced fungal diversity and evenness, while bacterial communities exhibited greater dominance and loss of rare taxa. These findings support the idea that fire promotes microbial post-fire niche specialization while disrupting dispersal pathways. Our results indicate that increasing fire frequency and severity under climate change may homogenize soil microbial communities, reduce microbial resilience, and constrain ecosystem recovery.

## Introduction

Fire is a global process that alters ecosystems and their biological communities [1,2]. Global changes are altering fire regimes worldwide [3] with negative consequences on global biological diversity [1], further threatening soil microbial diversity and soil biogeochemical cycles [4–7]. While previous studies showed that fire alters the composition and function of the soil microbiome [8,9], we still lack a general understanding of the effect of fire on the ecological processes driving the assembly of soil microbial communities, and how this can impact soil microbial diversity. These knowledge gaps constrain our ability to predict how varying fire regimes impact soil microbiome diversity and function, both cornerstones of global one health [10].

Fire produces instantaneous changes in soil microbial communities that persist after the passage of fire. Fire favors heat resistant or tolerant taxa (i.e., pyrophilic taxa), often selecting for species that are capable of either surviving fire itself, or in post-fire environments [11–13], and acts as a strong ecological selective force on soil microbial communities [14,15]. Yet, studies vary in predictions for how fire alters microbial communities [16,17]. Microbial community composition is linked to microbial functions like decomposition [18–20], fire-fuel feedbacks [21,22], and productivity [23]. Thus, understanding how fire influences microbial community assembly and diversity is imperative to understanding microbial roles in changing fire regimes.

The pyrodiversity-biodiversity hypothesis refers to fire’s ability to modify biodiversity through changes in ecosystem heterogeneity [24,25]. Generally, as fire intensity and severity increase, fire has a more homogeneous effect on soil conditions (e.g., C, N, P, pH, oxidative stress) [26–28] that leads to decreases in niche diversity and microbial biodiversity [25,29]. However, when fires are less intense, local variation in fire severity effects on soil conditions will produce a more heterogeneous environment that favors greater microbial β-diversity and stochasticity. While pyrodiversity-biodiversity relationships have been observed in other types of biological communities (e.g., plants) [30,31], this framework is seldom applied to soil microbial communities, representing another key knowledge gap in our understanding of fire-microbe interactions.

Fire likely impacts microbial stress tolerance and prevalence of specialists. Fire effects on ecosystems cause nutrient flushes [32,33], increases in stress (e.g., reduced soil water content, increased reactive oxygen species [34–36]), and changes in soil carbon (e.g., pyrogenic organic matter; PyOM) [37]. These novel post-fire conditions could alter the relative importance of generalist taxa in favor of more specialized microbes capable of surviving in burned ecosystems. For example, thermotolerant, fast growing taxa (e.g., Firmicutes, *Blastococcus*, and *Pedobacter*) often proliferate in burned soils [13,38,39] likely due to nutrient flushes, and PyOM specialists such as *Pholiota carbonaria* often proliferate on burned plant material after fire [40]. Thus, individual studies provide hints of how post-fire microbial communities shift, but a broader analysis is needed to identify rules for how fire influences microbial community assembly.

Fire could also influence microbial dispersal, which would alter microbial composition and function. Fire drives significant mortality in soil microbial communities that can reduce microbial biomass and competition [12,41]. This shift in community composition could favor ruderal taxa capable of dispersing into burned environments where they can take advantage of reduced competition and post-fire nutrient flushes [11,42]. Alternatively, fire produces harsh effects on ecosystems like soil drying, greater reactive oxygen stress, and the loss of plant hosts [35,36]. These less beneficial conditions may favor post-fire dispersal out of burned systems as microbes seek better conditions. This may explain the apparent increase in post-fire fruiting body formation [40,43], which could allow less fire tolerant taxa to escape post-burn environments. While a few studies have assessed microbial dispersal into and out of burned environments [44,45], the importance of post-fire dispersal to microbial community assembly has been largely ignored. Thus, there is a knowledge gap regarding our ability to predict the impact of increased fires with climate change on soil microbial community assembly via dispersal.

To address the key knowledge gaps identified above related to the effect of fire on the soil microbiomes, we used public amplicon sequencing data (>2,600 samples from 19 studies) to test three primary questions: 1) Does fire alter microbial community assembly processes, particularly ecological selection and dispersal? 2) Does fire influence microbial community diversity? 3) Does fire select for pyrophilic specialists and promote dispersal into burned sites? We hypothesized that fire would favor specialist taxa capable of surviving fire and post-burn ecosystems, driving homogenous selection on soil microbial communities. We also hypothesized that while fire would be associated with decreases in bacterial and fungal α-diversity, it would promote greater β-diversity and stochasticity (per the pyrodiversity-biodiversity hypothesis). Finally, we hypothesized that post-fire dispersal would be higher into burned environments given expected reductions in microbial diversity and biomass.

## Results

To test our hypotheses, we collected publicly available data from studies investigating the influence of fire on the soil microbiome using amplicon sequencing and targeting bacterial (16S rRNA gene) and fungal (ITS2 rRNA region) communities. Our first search resulted in 4,137 articles, which was then refined into 115 studies with a design relevant to answer our questions. This set was further reduced to 19 studies (16 for bacteria, 10 for fungi, of which 7 analyzed both communities), including only those studies with freely accessible data and compatible marker genes (i.e., located on the same region). Within each study, samples were further filtered, yielding 1,596 (bacteria) and 1,061 (fungi) high-quality samples.

We first tested whether fire alters the ecological processes behind soil microbiome assembly and, contrary to our hypothesis, we found that ecological selection processes were not altered by fire in bacterial and fungal communities (Fig. 1, Tab. S1). On the other hand, fire reduced the contribution of homogenizing dispersal in bacterial communities (Fig. 1A, Tab. S1). Similarly, we observed a reduction of homogenizing dispersal and dispersal limitation post-fire, together with an increase in the contribution of ecological drift (stochasticity) in the assembly of soil fungal communities (Fig. 1B, Tab. S1).

**Figure 1.**
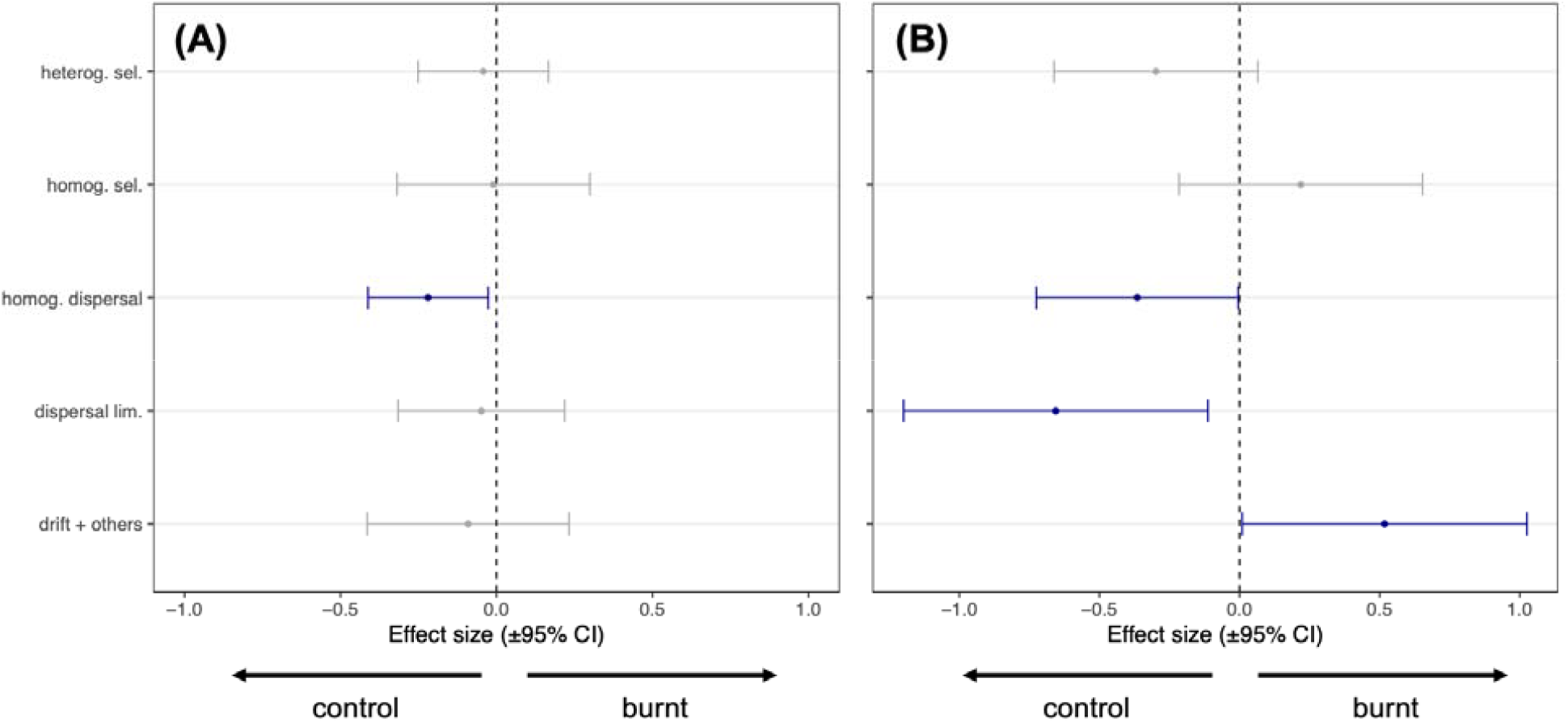
Effect size (±95% CI) of fire compared to unburned (control) sites on the ecological processes driving the assembly of soil (A) bacterial and (B) fungal communities. Bars in blue represent variables with effect size significantly different from zero (Tab. S1).

We predicted that fire would alter the niche partitioning between generalists and specialists, with an increased proportion of specialists after fire. Following our hypothesis, we indeed observed a higher proportion of ASVs identified as specialists in both bacteria (p = 0.02; Fig. 2A, Tab. S2-S3) and fungi (p = 0.001; Fig. 2B, Tab. S2-S3).

**Figure 2.**
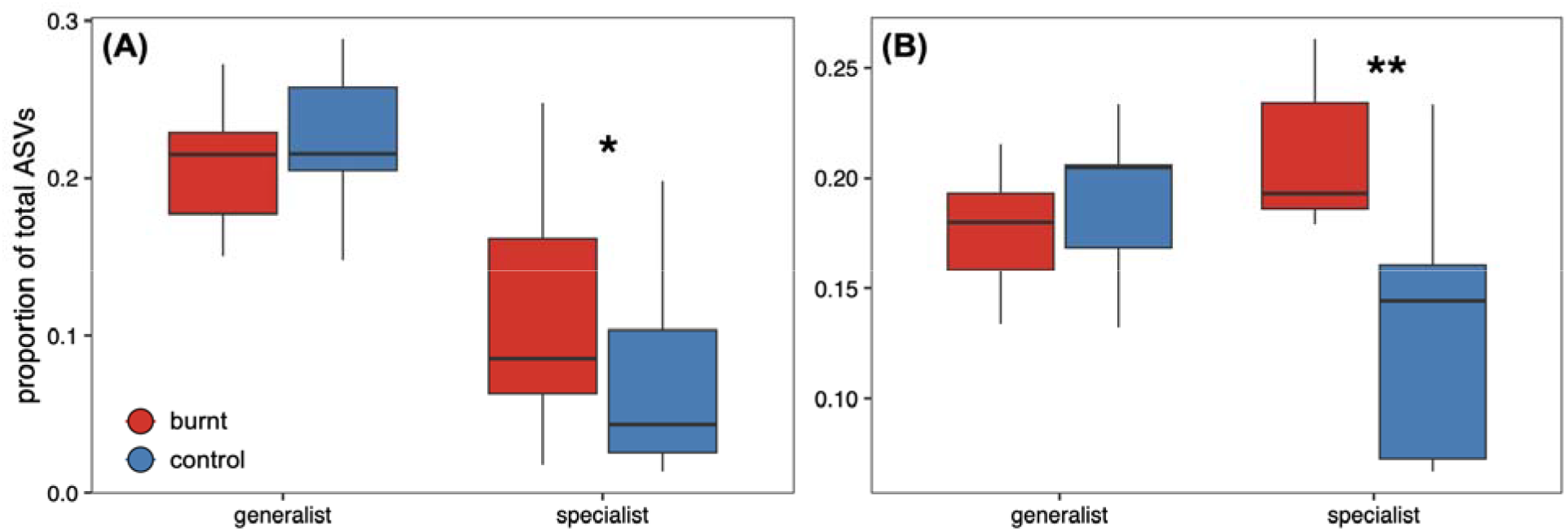
Effect size (±95% CI) of fire (red) compared to unburned (control, blue) sites on the proportion of specialists and generalists within soil (A) bacterial and (B) fungal communities. See Table S2-S3 for details on the model and post-hoc contrasts.

We also hypothesized that overall microbial diversity would be reduced in post-fire soil microbiomes. This proved true for fungi but not bacteria. In bacterial communities, we did observe no variation in any metric of microbial diversity (number of observed ASVs, Shannon diversity index, Chao 1 index, Faith’s phylogenetic diversity index; Fig. 3A, Tab. S1), but we observed an increase in dominance and a reduction in evenness and proportion of rare species (Fig. 3A, Tab. S1). On the other hand, in fungi, we observed an overall reduction in diversity without any variation in dominance and evenness (Fig. 3B, Tab. S1). We observed no variation in β-diversity (using the MNTD index) in either bacterial or fungal communities in post-fire soils (Fig. 3B, Tab. S1).

**Figure 3.**
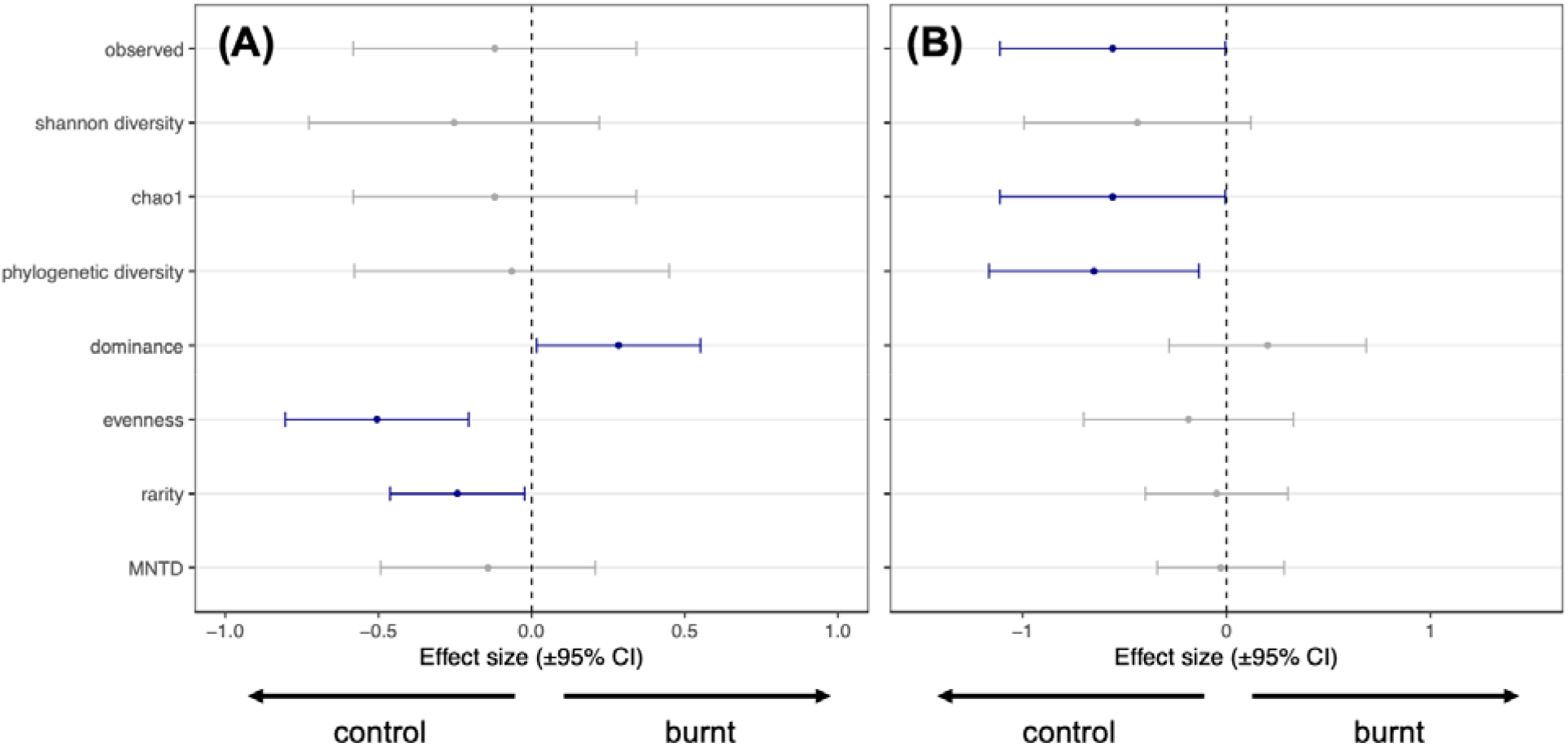
Effect size (±95% CI) of fire compared to unburned (control) sites on the diversity of soil (A) bacterial and (B) fungal communities. Bars in blue represent variables with effect size significantly different from zero (Tab. S1).

## Discussion

Our global analysis demonstrated that fire alters soil microbial community assembly patterns and limited dispersal, but these effects varied between bacteria and fungi. Bacteria show changes in dominance, evenness, and rarity, while fungi show overall decreases in diversity. While fire did drive clear changes in community assembly patterns, contrary to our expectations, there were no clear signals of homogenous or heterogenous selection on microbial communities. Further, fire effects on microbial diversity also varied between bacteria and fungi, with burned fungal communities becoming more heterogeneous after fire. This matches the expectations of the pyrodiversity-biodiversity hypothesis in that fire effects on ecosystem heterogeneity can produce more stochastic fungal communities where ecological drift plays a larger role in community assembly. Finally, fire also limited microbial dispersal into burned environments. This was contrary to our hypothesis, as we expected that post-fire reductions in diversity would allow for reduced competition and easier dispersal into burned systems.

Testing the effect of fire on soil microbial communities across fire types (e.g., wildfire and prescribed fire) and ecosystems allowed for generalizations about post-fire microbial community assembly rules. While some individual studies have found fire drives heterogeneous ecological selection [14,15], this effect was not apparent when generalizing across studies. Instead, we found that fire filters for pyrophilic specialists. This matches prior work reporting greater abundance of pyrophilic groups like Firmicutes [17,39] (heat resistant), Actinobacteria [13,38,46] (alkaline soil tolerant), *Pyronema* [47], and *Morchella* [48,49]. Fire produces a unique set of conditions (e.g., increased PyOM, reduced soil water content, increased soil pH, and nutrient flushes) which favor pyrophilic specialist microbes [35,36]. While many of fire’s effects on soils persist after fire, these effects do vary over time and may allow for unique post-fire microbial successional trajectories [39,50–52]. In summary, fire associated changes in microbial community assembly reflect filtering for pyrophilic specialists.

While fire selected for pyrophilic specialists, the mechanisms for this selection varied between soil bacteria and fungi. For bacteria, fire promoted the dominance of key taxa while reducing community evenness and the number of rare species rather than reducing diversity. This suggests that bacteria killed by fire are quickly replaced by pyrophilic specialists (i.e., species turnover), and reflects the findings of previous studies which have demonstrated that post-fire bacterial community assembly is more deterministic [53] and linked to selection for heat resistance and adaptation to burned soils [54,55]. Relative to bacteria, fire effects on fungi were more associated with mortality (i.e., lower diversity in burned sites). This is supported by past studies reporting that fungal diversity tends to be more adversely affected by fire relative to bacteria [13,41,56]. This effect may be driven by longer fungal generation times, the loss of plant hosts during fire, as well as larger, less thermotolerant cells [11]. Summarizing, while fire selected for pyrophilic specialists, the mechanisms varied between bacteria (species turnover) and fungi (mortality).

Fire effects on bacterial and fungal diversity varied, and with burned bacterial and fungal communities becoming more deterministic and stochastic respectively. While the mean nearest taxon distance (MNTD; phylogenetic β-diversity metric) did not vary between burned and unburned bacterial or fungal communities, other diversity metrics were strongly influenced by fire. Amongst bacteria, fire promoted communities with fewer rare taxa that were less even and favored the dominance of some taxa. While other studies have found that stochastic processes are important contributors to post-fire bacterial community assembly [14,29], we did not find a strong overall signal of homogenous or heterogeneous selection. Fungal communities however did become more stochastic after fire (greater contribution of ecological drift) even though fire generally reduced fungal diversity. This provides varying levels of support for the pyrodiversity-biodiversity hypothesis (i.e., greater ecosystem heterogeneity after fire promotes greater biodiversity [26–28]). While past work has shown that ecosystem heterogeneity introduced by fire promotes diverse microhabitats and local variation in soil microbial communities [29,57], here we only found support for this with soil fungi. The variation in findings may be due to differences in spatial scale and fire severity amongst the included studies [29,57], as pyrodiversity-biodiversity effects can be obscured if spatial sampling scales are too large or reduced if only higher severity wildfires are considered. While more work is required to determine the appropriate spatial and temporal scales for assessing pyrodiversity-biodiversity relationships between fire types and ecosystems, this demonstrates the ability of fire to introduce abiotic and biotic heterogeneity into ecosystems. If fire-driven increases in microbial diversity also influence plant community composition and productivity [58], then it is likely that pyrodiversity-biodiversity relationships underlie the fire-fuel feedbacks that promote recurrent fire in pyrophilic grasslands, savannas, and forests [59–61].

Fire reduced dispersal into post-fire soil microbial communities. While fire did drive reductions in microbial diversity, potentially reducing interspecies competition, this did not coincide with increased microbial dispersal into burned sites. Although past studies have found that post-fire dispersal can contribute to airborne microbial propagules and influence microbial community assembly, this is restricted to litter microbial communities and varies with time since fire [44,45,62]. This suggests that airborne fungal spores found during and after fire are taxa that disperse out of burned ecosystems rather than into them [11,28] and that post-fire fungal fruiting bodies are evidence of dispersal out of burned sites [40,63,64]. Thus, taxa capable of taking advantage of post-fire nutrient flushes and soil conditions are likely already present in burned soil and not rapid dispersers that take advantage of post-fire nutrient flushes [11,28]. Reductions in dispersal into burned areas is likely to reduce recovery and extend the impact of fires on microbial function.

The impact of fire on microbial community assembly suggests climate change effects on fire regimes will drive lasting changes in soil microbial communities and their roles in ecosystems. Climate models predict that fire frequency and the amount of burned area will increase [3,65,66], therefore we expect decreases in soil bacterial and fungal α-diversity, and increases in fungal β-diversity driven by greater stochasticity and drift in community assembly. As fire conditions deviate from historical norms due to fuel build, fire suppression, and/or drought [67–69], higher severity fires will homogeneously drive mass microbial mortality, loss of diversity, and limit microbial functions (e.g., decomposition) [19,20,70,71]. Further, since fire limits microbial dispersal into burned ecosystems, post-fire microbial recovery will likely be slower as fire severity increases relative to historic norms. Given the expected changes in fire regimes, increasing the use of prescribed fire and management that encourages natural fire regimes will be critical to the conservation of soil microbial communities and ecosystems.

In conclusion, fire changes soil microbial community assembly patterns, promotes fungal pyrodiversity-biodiversity relationships, and limits dispersal into burned environments. Fire filtered for specialist pyrophilic microbial taxa, but the mechanisms varied between bacteria (e.g., species turnover, changes in dominance) and fungi (e.g., mortality and loss of diversity). While this meta analysis addresses major questions in microbial fire ecology, the lack of data in regard to the effect of fire type, fire frequency, fire severity, time since fire, and fire season (i.e., spring vs. fall) demonstrated key knowledge gaps going forward. The dataset presented here was biased towards North American forests and grasslands, and thus in the future there are opportunities to not only address how the fire regime components mentioned above influence soil microbial communities, but also test how fire effects on microbes vary across ecosystem types and regions. The importance of understanding soil microbial responses to fire is only expected to increase as the effects of climate change become more apparent.

## Methods

Data collection was performed using a similar approach to Malacrinò et al. [72]. The search for the term “fire AND micro” on the Web of Science Core Collection (year range 2006–2024) yielded 4,137 records (accessed on July 3, 2024). To guarantee completeness of the data set, a Google scholar search using the keywords: “fire AND illumina AND microb* OR fung* OR bact* or archae*” was used (added 43 records). We then filtered this first dataset to remove all those papers not performing amplicon sequencing on soil samples in post-fire conditions.

Our aim was to develop a very robust analysis, so we took additional steps rarely performed in meta-analyses of sequencing data. We additionally filtered this set of studies focusing on those: (i) with publicly available and accessible data deposited in SRA databases; (ii) that used Illumina sequencing technology to sequence the V3–V4 portion of the 16S rRNA gene (bacterial community) and/or the ITS2 rRNA region (fungal community). This allowed us to design a replicable pipeline and reduce the bias generated by comparing data obtained from different sequencing platforms or from sequencing different regions of marker genes. Studies that did not specify the primer pairs used, lacked clear differentiation between samples when multiple markers were employed, or contained non-demultiplexed data, were excluded from the analysis. This selection process resulted in 28 studies. Within each study, samples underwent additional filtering (see below), resulting in 19 studies included in our meta-analysis (16 focusing on bacteria, 10 on fungi, with 7 examining both communities) with a total of 1,596 high-quality samples for bacteria and 1,061 for fungi. To facilitate information tracking, each study and sample was assigned a unique ID number: the study ID was the BioProject number, and the sample ID was the SRA sample number. For each sample, the metadata we collected included geographic location and site type (burnt, control). Data were subsequently downloaded using SRA Toolkit 3.0.2 (see Tab. S4 for the list of studies and samples included in our analyses).

Raw data was carefully handled to reduce study-specific bias. Although we selected studies that characterized the same region within each marker gene for amplicon sequencing, there were slight inconsistencies in the primer pair used for PCR amplifications. To address this possible source of bias, we generated a custom script using *Cutadapt* v5.0 to remove primer sequences from each individual sample using the information available from each study. During this step we also removed low quality reads and very short reads using *Cutadapt* default settings. Reads were then processed with the *nf-core/ampliseq* v2.10.0 pipeline [73–75] for additional quality control, error correction, clustering of ASVs, and chimera removal. ASV sequences were aligned using *MAFFT* v7.525 [76] and used to build a phylogenetic tree using *FastTree* v2.1.11 [77]. Taxonomic assignment was performed using SILVA database v138.1 [78] for the 16S dataset, and the UNITE database v10.0 [79] for the ITS dataset. All subsequent data processing and analysis were performed in *R* v4.4.1 [80], after merging the ASV table, taxonomic identification table, metadata, and phylogenetic tree using *phyloseq* v1.48 [81]. Singletons, ASVs identified as mitochondria or chloroplasts, and samples with less than 1,000 reads were discarded from the dataset.

The following diversity metrics were calculated using the package *microbiome* v1.26 (https://github.com/microbiome/microbiome): observed number of ASVs, Chao1 index, Shannon diversity index, Berger–Parker dominance index, Pielou evenness index, Log-Modulo Skewness rarity index. The Faith’s phylogenetic diversity and the MNTD (Mean Nearest Taxon Diversity) were calculated using the package *picante* v1.8.2 [82]. The relative contribution of different ecological processes in the assembly of the soil bacterial and fungal microbiomes was estimated using the package *iCAMP* v1.5.12 [83]. Each ASV was classified into “specialist” or “generalist” using the package *EcolUtils* v0.1 (https://github.com/GuillemSalazar/EcolUtils). Diversity metrics and the relative contribution of each assembly process were used to calculate the mean, standard deviation, and sample size for each study and sample type (burnt vs control), and then this dataset was used to calculate Standard Mean Difference using the package *metafor* v4.8 [84]. This data was then used to fit a meta-regression linear mixed-effects model using the package *metafor*. The proportion of ASVs identified as “specialist” or “generalist” was instead fit to a linear mixed-effect model using the package *lme4* v1.1 [85] using the variable sample type (burnt, control), niche (specialist, generalist), and their interactions as fixed factors, and the variable “study ID” as random effect, while post-hoc contrasts were extracted using the package *emmeans* v1.10.7 [86].

## Supporting information

Supplementary materia

## Acknowledgements

AM was supported by the Next Generation EU - Italian NRRP, Mission 4, Component 2, Investment 1.5, call for the creation and strengthening of ‘Innovation Ecosystems’, building ‘Territorial R&D Leaders’ (Directorial Decree n. 2021/3277) - project Tech4You - Technologies for climate change adaptation and quality of life improvement, n. ECS0000009. This work reflects only the authors’ views and opinions, neither the Ministry for University and Research nor the European Commission can be considered responsible for them. We acknowledge the support by Clemson University’s HPC resources [87]. AEB was supported by The Ohio State University. JRH was supported through The Ohio State University’s President’s Postdoctoral Scholar Program.

## Author contribution

Conceptualization: JRH, AM, AEB; Methodology: JRH, AM; Investigation: JRH, GV, AM; Visualization: AM; Writing-original draft: JRH, AM; Writing-review and editing: all co-authors.

## Data and code availability

Raw data from individual studies are publicly available on NCBI SRA (see Supplementary Table S4). The code used to perform analyses is available on GitHub: https://github.com/amalacrino/fire_meta_analysis

