## Supplementary materia for "Fire limits soil microbial dispersal and differentially impacts bacterial and fungal communities"

### Supplementary material

**Table S1**. Results from the meta-regression models testing the influence of fire (burned vs control) on the ecological processes of microbiome assembly and diversity metrics for bacterial and fungal communities.

| **Metric** | **Bacterial community** | | **Fungal community** | |
|---|---|---|---|---|
| | $Q_M$ | **p** | $Q_M$ | **p** |
| **Heterogeneous selection** | 0.15 | 0.68 | 2.57 | 0.11 |
| **Homogenous selection** | 0.01 | 0.95 | 0.97 | 0.32 |
| **Dispersal limitation** | 0.12 | 0.72 | 5.61 | 0.01 |
| **Homogenizing dispersal** | 4.98 | 0.02 | 3.94 | 0.04 |
| **Drift and others** | 0.30 | 0.58 | 3.98 | 0.04 |
| **Observed number of ASVs** | 0.25 | 0.61 | 3.92 | 0.04 |
| **Chao 1 index** | 0.26 | 0.61 | 3.92 | 0.04 |
| **Shannon diversity index** | 1.09 | 0.29 | 2.36 | 0.12 |
| **Dominance (Berger–Parker index)** | 4.31 | 0.03 | 0.66 | 0.41 |
| **Evenness (Pielou)** | 10.92 | 0.0009 | 0.51 | 0.47 |
| **Rarity index (Log-Modulo Skewness)** | 4.68 | 0.03 | 0.07 | 0.78 |
| **Faith's phylogenetic diversity index** | 0.06 | 0.81 | 6.12 | 0.01 |
| **Mean Nearest Taxon Diversity (MNTD)** | 0.63 | 0.42 | 0.03 | 0.85 |

**Table S2**. Results from the linear mixed-effects model testing the influence of fire (burned vs control) on the proportion of specialists and generalists within bacterial and fungal communities.

| | **Bacterial community** | | | **Fungal community** | | |
|---|---|---|---|---|---|---|
| | $\chi 2$ | **df** | **p** | $\chi 2$ | **df** | **p** |
| **fire** | 1.21 | 1 | 0.27 | 3.47 | 1 | 0.06 |
| **niche group** | 96.01 | 1 | < 0.001 | 1.56 | 1 | 0.21 |
| **fire × niche group** | 5.26 | 1 | 0.02 | 10.32 | 1 | 0.001 |

**Table S3**. Post-hoc contrasts (burned vs control) of proportion of specialists and generalists within bacterial and fungal communities.

| | **Bacterial community** | | **Fungal community** | |
|---|---|---|---|---|
| | **estimate** | **p** | **estimate** | **p** |
| **generalists** | -0.01 | 0.41 | -0.01 | 0.35 |
| **specialists** | 0.04 | 0.02 | 0.06 | 0.001 |

**Table S4**. Studies included in this meta-analysis. See XLSX file.
